# Selection of Extended CRISPR RNAs with Enhanced Targeting and Specificity

**DOI:** 10.1101/2023.01.11.523593

**Authors:** Ashley Herring-Nicholas, Hillary Dimig, Miranda Roesing, Eric A. Josephs

## Abstract

For a CRISPR guide RNA (gRNA) with a specific target but activity at known “off-target” sequences, we present a method to screen hundreds of thousands of gRNA variants with short, randomized 5’ nucleotide extensions near its DNA-targeting segment—a modification that can increase Cas9 gene editing specificity by orders of magnitude with certain 5’- extension sequences, *via* some as-yet-unknown mechanism that makes *de novo* design of the extension sequence difficult to perform manually—to robustly identify extended gRNAs (x-gRNAs) that have been counter-selected against activity at those off-target sites and that exhibit significantly enhanced Cas9 specificity for their intended targets.

## MAIN TEXT

CRISPR effector Cas9 from *Streptococcus pyogenes* (SpyCas9) has emerged over the past several years as a powerful biotechnological tool that also holds tremendous therapeutic potential in the treatment of genetic diseases.^1,2^ This potential arises from the ability of CRISPR effectors to use a modular segment of their RNA co-factor, their guide RNA or ‘gRNA’, to recognize DNA sequences complementary to its ‘spacer’ segment and introduce targeted mutations into the DNA at those sites.^3,4^ However, oftentimes a gRNA for a specific target can cause the Cas9 nuclease to introduce “off-target” double-strand breaks (DSBs) and mutations at similar nucleotide sequences that are also present in that genome,^5,6^ and the possibility of unintended or uncontrolled Cas9-induced mutational events raises significant concerns for those therapeutic applications. In particular, it is becoming increasingly important to recognize that individuals may carry unique or “personal” off-target sequences for a therapeutic gRNA as a result of genetic variations that exist between people and/or across different populations, and that these unique off-target sequences must be accounted for in an era of personalized medicine.^7,8^

Although there have not been any approaches developed to directly limit activity at those specifically-identified off-target sequences for a gRNA of interest, there are a few ways to reduce “off-target” activity overall and increase the specificity of CRISPR systems in general. For example, these general approaches include reducing cellular exposure to Cas9 nucleases^9^ or selectively inhibiting Cas9 nuclease activity altogether,^10^ as well as using engineered, “high fidelity” or “enhanced specificity” Cas9 variants such as eCas9.^11^ eCas9 effectors have amino acid substitutions designed to reduce their overall affinity for DNA in a way that decreases the probability that the effectors’ latent nuclease domains will become activated at sequences with imperfect complementarity to its gRNA.^12^ Modification of the gRNA itself has also been found to modulate Cas9 specificity: gRNAs with chemically modified bases, phosphates, or sugars can exhibit increased specificity overall relative to unmodified gRNAs, although the optimal combination of modifications for a specific target/off-target can be difficult to predict *de novo*.^13,14^ Removing a few nucleotides from the 5’ end of the DNA-targeting segment of the gRNA (from 20 nt to 17 or 18 nt) to generate truncated gRNA (‘tru-gRNAs’) can also decrease off-target activity,^15^ an effect likely caused by a general destabilization of gRNA/DNA interactions when the spacers are shortened.^12^

Recently, it was found that adding short nucleotide extensions (∼6 to ∼16 nts) to the 5’- end of the gRNA next to its DNA-targeting ‘spacer’ segment (Figure 1B)—especially those that were predicted to form ‘hairpin’ or secondary structures with the spacer designed to interfere with gRNA interactions at specific off-target sequences—could significantly reduce Cas9 off-target activity while maintaining on-target mutational efficiencies.^16^ On average, the specificity in targeting during gene editing for these “hairpin-gRNAs” or “hp-gRNAs” increased 50-fold (and up to 200-fold) relative to gene editing using standard gRNAs, and this approach worked in diverse CRISPR effectors for multiple target sites each. While those hp-gRNAs could significantly outperform the state-of-the-art, the 5’ extended sequences were each designed and tested one-at-a-time, manually, for each targeted sequence and set of off-targets. At the time, it was also found that some of tested 5’ - extensions did not effectively reduce off-target activity; others also significantly inhibited on-target activity; and still others (controls) that were not predicted to increase specificity occasionally did. Because there are a very large number of possible short 5’-extensions and because, in principle, different 5’-extensions for the same spacer sequence can be fine-tuned or optimized to limit activity *vs*. specific off-target sequences, the inability to predict *de novo* which of those sequences will increase the specificity of an associated Cas9/gRNA ribonucleoprotein (RNP) has so far limited their utility in practice in eliminating the risk of off-target mutation during gene editing.

**Figure 1.**
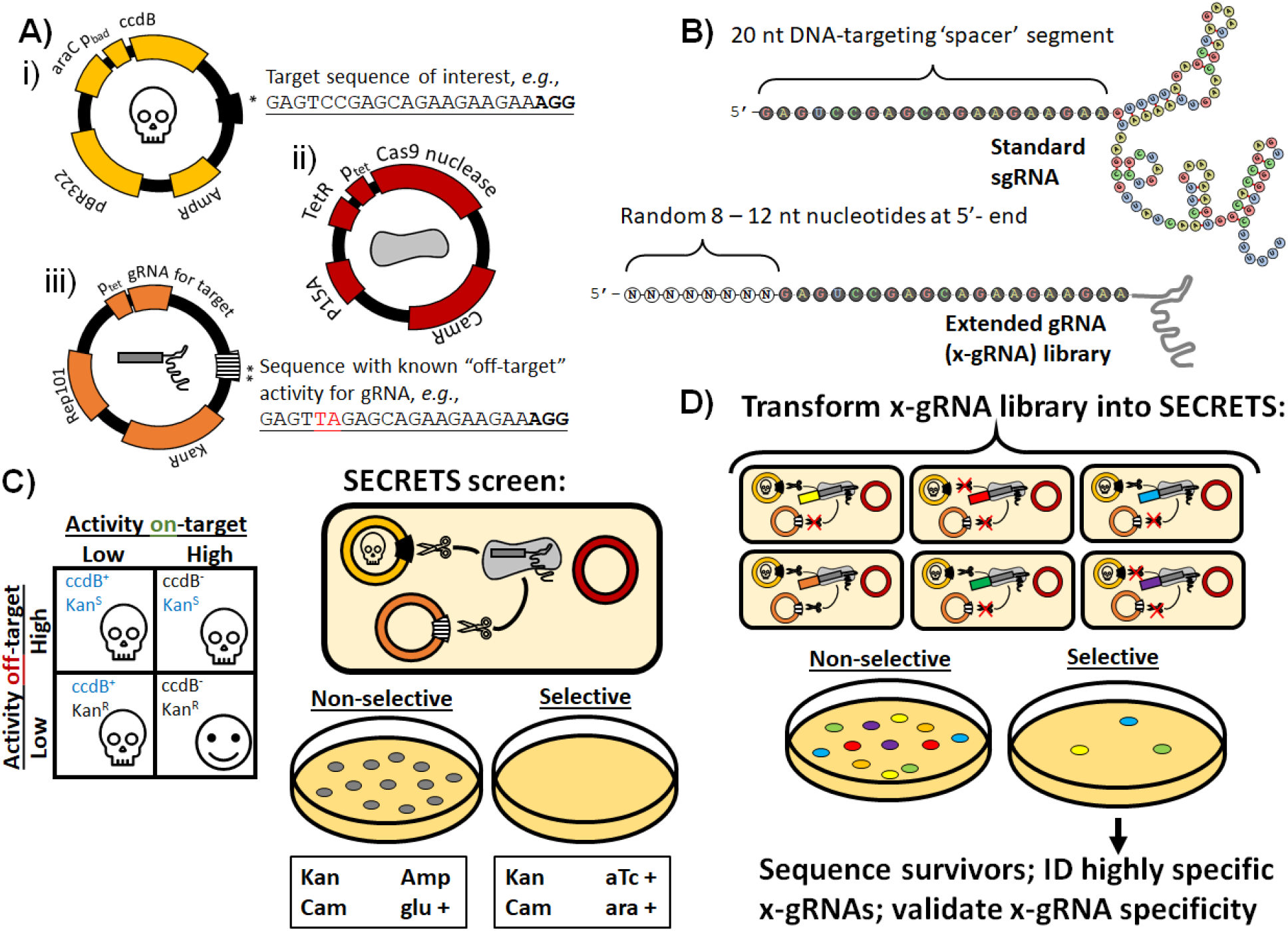
SECRETS screen to identify short 5’-nucleotide extensions to gRNAs that increase Cas9 gene editing specificity for a specific target with known off-targets. A) Simplified schematics of the three SECRETS plasmids: (i) high-copy number plasmid with inducible toxin and target sequence of interest to select for “on-target” activity, (ii) medium copy plasmid for inducible expression of Cas9, and (iii) low-copy plasmid for gRNA expression and counterselection for “off-target” activity with kanamycin resistance cassette and a known “off-target” sequence for the gRNA. B) (above) a standard gRNA has 20 nt “spacer” (DNA-targeting) segment while (below) an extended gRNA (x-gRNA) has an additional 8 – 12 nt to the 5’-of the spacer. C) During SECRETS selection, *E. coli* expressing standard gRNAs that exhibit promiscuous activity are unable to survive. D) Members of large, randomized x-gRNA libraries can be screened simultaneously using SECRETS for 5’-extensions that result in efficient on-target Cas9 activity and minimized off-target activity that can be used for highly-specific gene editing applications at that target.

To overcome this challenge, here we present an experimental protocol to screen tens-to hundreds-of thousands of candidate 5’-extension sequences simultaneously to efficiently and reliably identify novel to extended gRNA sequences (x-gRNAs) that maintain robust Cas9 activity on-target while significantly increasing their gene editing specificity bu effectively eliminating their activity at known off-target sequences where conventional approaches to increase Cas9 specificity in general may fail (Figure 1). In this protocol, called “<u>S</u>election of <u>E</u>xtended <u>C</u>RISPR <u>R</u>NAs with <u>E</u>nhanced <u>T</u>argeting and <u>S</u>pecificity” (SECRETS), the activities of Cas9 enzymes with a library of x-gRNA candidates are evaluated in parallel using an *Escherichia coli* strain that is strongly selective for their ability to stimulate Cas9 nuclease activity on-target and strongly counter-selective for activity at their off-targets. The SECRETS *E. coli* strain maintains three plasmids (Figure 1A): (i) a high-copy plasmid expresses the toxin ccdB in the presence of arabinose (ara), and also contains the target sequence of interest; (ii) a medium-copy plasmid expresses Cas9 in the presence of anhydrotetracycline (aTc); and (iii) a low-copy plasmid expresses the gRNA of interest, provides resistance to the antibiotic kanamycin (KanR), and also contains a known off-target sequence for the gRNA. DSBs induced by the Cas9 at the target of interest *in E. coli* results in the degradation of ccdB plasmid, allowing the bacteria to survive in the presence of arabinose, while DSBs induced by the Cas9 at the known off-target results in the degradation of the gRNA plasmid and bacterial susceptibility to the antibiotic kanamycin (KanS).^17^ Hence, only gRNAs that exhibit robust activity at their intended targets (degrading the high-copy toxin plasmid) and low activity at their off-target sites (not degrading the low-copy gRNA plasmid) will survive a SECRETS screen (Figure 1C).^18^

We tested this system with a well-characterized gRNA^16^ for human gene *EMX1*, its target sequence (*EMX1* ON), and another sequence found in the human genome where the Cas9/gRNA RNP complex is known to exhibit off-target activity (*EMX1* OFF1) due to a two nucleotide difference with the *EMX1* ON target at positions where Cas9 effectors are especially susceptible to tolerating sequence divergence (Figure 1A).^19^ After only 1 hr of Cas9/gRNA expression, followed by plating and overnight growth on LB with aTc, arabinose, chloramphenicol (cam), and kanamycin, we find strong suppression of *E. coli* growth with the standard *EMX1* gRNA, but only when the *EMX1* OFF1 sequence is present in the kanamycin resistance plasmid (Figure S1). If instead of the standard gRNA for EMX1 (Figure 1B top) we introduce a library of EMX1 x-gRNA variants with 8 randomized nucleotides (N8) appended to its 5’ end (Figure 1B bottom), we find numerous *E. coli* colonies of survivors of the SECRETS protocol (Figures 1D and S1), indicating that these x-gRNAs from the library demonstrate the high Cas9 activity and specificity required to survive. The pooled survivors were sequenced and each of the top five most prevalent x-gRNA sequences in the surviving population (Figure 2A) were tested for activity and specificity *in vitro* (Figure 2B and S2, Table S1). *In vitro* Cas9 digestion assays revealed nuclease activities of Cas9 with all five of the x-gRNAs identified from the SECRETS screen had significantly reduced off-target activity at *EXM1* OFF1 compared to the standard *EMX1* gRNA—effectively eliminating nuclease activity at the known off-target site—and exhibiting similar activity at *EMX1* ON as the engineered “enhanced specificity” Cas9 variant eCas9. These five x-gRNAs identified through the SECRETS protocol could also exhibit higher levels of specificity in general, eliminating off-target activity across three other known *EMX1* off-targets (*EMX1* OFF2 – OFF4, containing 2 to 4 differences with the *EMX1* ON sequence) for Cas9, and reducing nuclease activity at all four off-target sequences even more so relative to eCas9 with a standard gRNA (Figure 2B and S2). In addition to x-gRNAs for *EMX1*, we were also able to readily identify multiple x-gRNAs using the SECRETS protocol for other targets—human genes *VEGFA* and *FANCF* (Figure S3 and Table S1)^16^— with superior activity and specificity profiles (Figure 2C), including in cases (*VEGFA*) where eCas9 was not able to significantly reduce activity at OFF1 sites.

**Figure 2.**
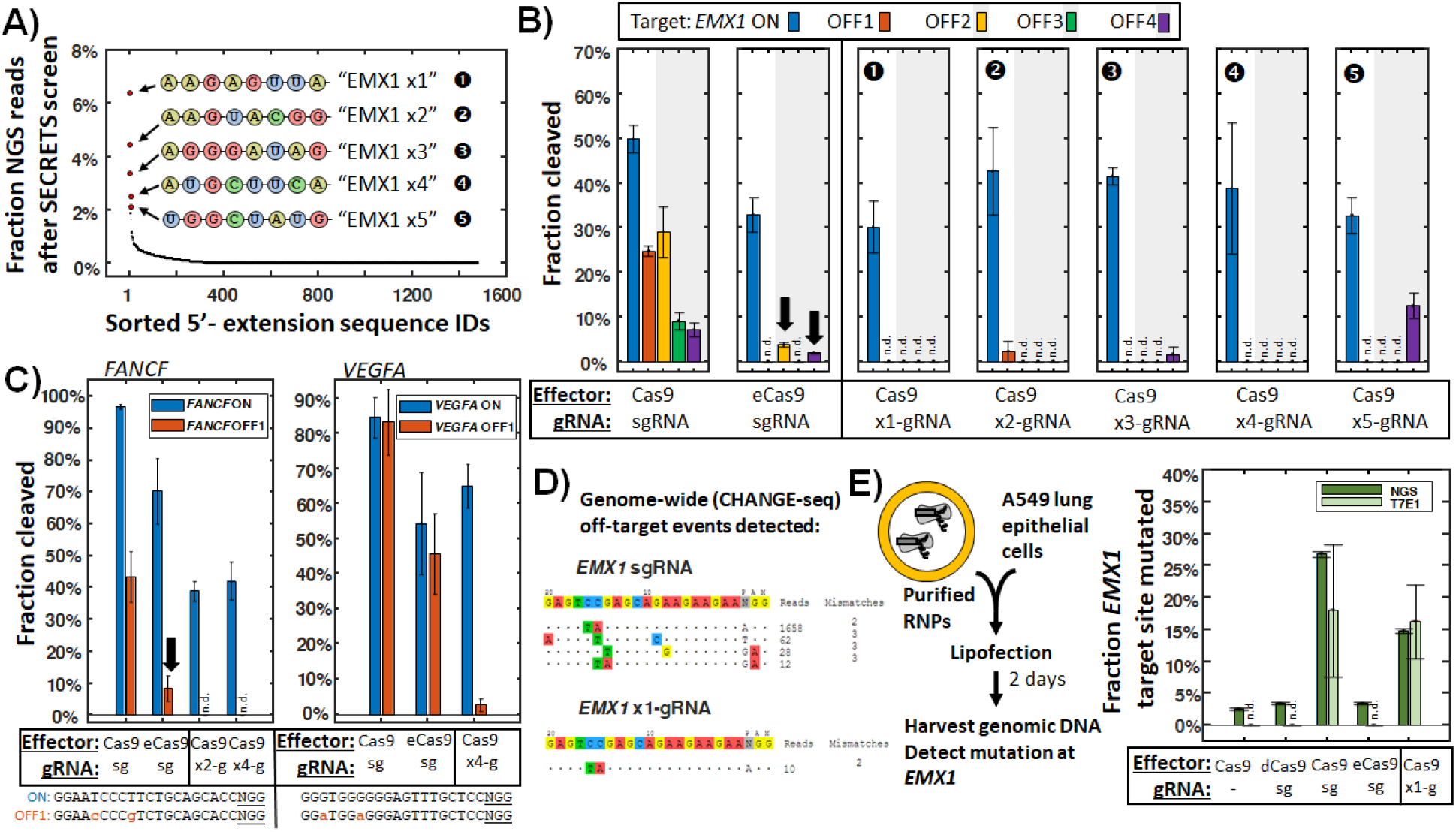
Selection and validation of x-gRNAs. A) The top five x-gRNAs recovered from two SECRETS screen replicates for the *EMX1* gRNA were identified *via* next-generation sequencing (NGS) for *in vitro* validation. B) All five x-gRNAs exhibit remarkable on-target activity (comparable or greater than “enhanced specificity” Cas9 variant eCas9) and greatly reduced off-target activities across four known off-target sites for the standard x-gRNA during in vitro cleavage assays using purified Cas9 and *in vitro*-transcribed (x-)gRNAs. n = 3 replicates each. C) Highly-specific x-gRNAs could also be readily identified from SECRETS screens for gRNAs for *FANCF* and *VEGFA* targets. n = 3 replicates each. D) CHANGE-seq^5^ reveals that x-gRNAs identified through SECRETS exhibit minimal and greatly-reduced genome-wide off-target activity, with no novel off-target activity identified relative to the standard gRNA. n = 2 biological replicates each. E) x-gRNAs identified through SECRETS exhibit robust gene editing activity following transfection of purified Cas9 ribonucleoprotein (RNP) complexes. sg = standard gRNA, dCas9 = catalytically inactive Cas9, eCas9 = enhanced specificity Cas9, NGS = next-generation (illumina) sequencing, T7E1 = T7E1 mutation detection assay. n = 3 biological replicates, 2 NGS replicates, and 3 T7E1 replicates.

To confirm that the x-gRNAs identified *via* SECRETS would remain active in human cell lines for gene editing, we transfected RNP complexes with Cas9 variants (wild-type Cas9; catalytically-inactive dCas9;^4^ or engineered enhanced-specificity eCas9^11^) and gRNA variants (standard gRNAs or x-gRNAs) into A549 human lung epithelial cells, then performed T7E1 mutation detection assays and next-generation sequencing (NGS) to quantify mutation rates at the *EMX1* target. The Cas9 RNPs with x-gRNAs identified *via* SECRETS exhibited robust gene editing cells in A549 cells, and higher on-target mutation rates than eCas9 (Figures 2E). We note that, even though they have additional 5’-nucleotides, hp-gRNAs had previously been found not to recognize or mutate any novel off-target sites compared to the standard gRNAs,^16^ likely because the 20 nt targeting segment of the hp-gRNA remains the same. To ensure that no new off-targets were introduced when using x-gRNAs identified from the SECRETS screen we also performed a test of genome-wide off-target nuclease activity (CHANGE-seq^5^) and, as expected, we found significant reductions of off-target cleavage activity genome-wide and no new off-targets (Figure 2D). Therefore, these findings demonstrate that the SECRETS protocol can robustly identify multiple high-performance x-gRNA candidates with strong potential for specific gene editing applications in human cells that eliminate off-target activity at selected loci.

In the demonstrations above, we screened randomized x-gRNA libraries of 65,336 variants (N8) in the SECRETS protocol for each gRNA target. We also identified enhanced x-gRNAs from more complex initial libraries (>250,000 5’-extension sequence variants), including pooled libraries of N8 variants containing different additional 4 nt tetraloop motifs^16^ (N8+4) designed to promote interactions between the N8 segment and the DNA-targeting segment of the x-gRNA (Figure S3). Indeed, we note that the ‘space’ of potential 5’-extension sequences for x-gRNAs is quite large^16^ (Figure S3)—we expect larger x-gRNA libraries of >4^10 or 4^12 (>1 - 16M) variants can be quite readily generated and screened in *E. coli* for new targets/off-target pairs of interest. The results here suggest that this space is also quite rich with high performance x-gRNA variants, as we could identify multiple x-gRNAs for several spacer sequences under relatively small-scale SECRETS screens that exhibited exceptional activity and specificity profiles. Furthermore, hp-gRNAs were able to improve the specificity of diverse CRISPR effectors, including Cas9 from *Staphylococcus aureus* and various Cas12 effectors,^16^ the SECRETS protocol can be readily adapted to those systems as well (Figure S3).

For many biotechnological or therapeutic applications, it is often desirable direct a Cas9/gRNA RNP to a specific nucleotide target of interest (*e*.*g*., where there might be no flexibility to target nearby sites). If it is determined there is a potential for off-target activity at certain sites in a specific sample or for a patient, there can be limited options to limit that activity at those specific off-targets. Here we demonstrate that the SECRETS protocol can be used to robustly identify ultra-specific variants for those gRNAs of interest that have been explicitly counter-selected against activity at those off-target sites. This approach could be used to robustly generate x-gRNAs in a “design-free” way that effectively eliminates the need for individualized optimization, and has been experimentally streamlined for simplicity in cloning new target/off-target pairs on-demand into the screening plasmids and for ease of rapidly selecting enhanced x-gRNAs. Once any off-target activity has been suspected or characterized, the SECRETS screen therefore provides an accessible and reliable method to identify high-performance gRNA variants for specific targets of interest. We expect the continued output and development of this approach will allow for safer applications of advanced CRISPR gene editing approaches that require gRNAs with extreme specificity, such as SNP-targeting and/or allele-specific gene editing. As approaches to predicting and identifying novel off-target sequences at the level of individual patients become more sophisticated and routine,^5,7^ we expect methods like SECRETS, which can reliably and rapidly generate highly-specific and highly-active gRNA variants that effectively eliminate Cas9 activity at specific off-targets, will become increasing important in applications of gene therapies in personalized medicine.

## Materials and Methods

### DNA oligonucleotides, dsDNA, and plasmids

DNA sequences for all oligonucleotides, dsDNA fragments, and plasmids are listed in Tables S2, S3, and S4, respectively.

### Cell lines and E. coli strains

All cloning was performed using New England Biolabs (NEB) 10-beta cells (NEB #C3020K) or TOP10 (Invitrogen #C404010) cells, and all SECRETS assays performed in Stbl2 cells (Invitrogen #10268019), grown at 30°C. A549 (ATCC CCL-185™) human lung epithelial cells were obtained from were obtained from ATCC (American Type Culture Collection).

### Cloning SECRETS plasmids and x-gRNA libraries

#### SECRETS plasmids

Three plasmids were generated for the validation of the SECRETS protocol: pSECRETS-A (medium copy p15A ori, chloramphenicol resistance, aTc-inducible Cas9 expression; Figure 1Aii), pSECRETS-B (low copy SC101 ori, kanamycin resistance, aTc-inducible gRNA expression, and a site for “off-target” sequence; Figure 1Aiii), and pSECRETS-C (p11.LacY.wtx1 plasmid (Addgene #69056) high copy pBR322 ori, ampicillin resistance, arabinose-inducible/glucose-suppressed ccdB toxin also containing additional site for “target” sequence; Figure 1Ai). p11-LacY-wtx1 was a gift from Huimin Zhao (Addgene plasmid # 69056 ; http://n2t.net/addgene:69056 ; RRID:Addgene_69056).

To clone pSECRETS-A, the Cas9 gene was PCR amplified from pwtCas9-bacteria (Addgene #44250) and a gBlock (purchased from Integrated DNA Technologies [IDT]) containing an anhydrotetracycline (aTc) inducible promoter (pLTetO-1) were inserted *via* HiFi Assembly (HiFi Assembly Kit (NEB#E5520S)) into a PCR-ed fragment of plasmid pBbA2c-RFP (Addgene #35326) to replace the red fluorescent protein (RFP) gene. pwtCas9-bacteria was a gift from Stanley Qi (Addgene plasmid # 44250 ; http://n2t.net/addgene:44250 ; RRID:Addgene_44250). pBbA2c-RFP was a gift from Jay Keasling (Addgene plasmid # 35326 ; http://n2t.net/addgene:35326 ; RRID:Addgene_35326).

To clone pSECRETS-B, the gRNA cassette for aTc-induced expression was constructed by inserting *via* HiFi Assembly a PCR’ed fragment of TetR from pBbA2c-RFP (Addgene #35326) and a gBlock (IDT) containing the pLTetO-1 promoter and a Golden Gate cassette (dual BsaI restriction sites) near a Cas9 fused tracrRNA-crRNA fusion to replace the RFP and LacI genes in a PCR’ed fragment of pBbS2K-RFP (Addgene #35330). The Golden Gate assembly cassettes of the resulting plasmid (called pSECRETS-B) could then be used to clone spacer sequences or x-gRNA libraries for gRNA expression after BsaI digestion and ligation of short phosphorylated annealed oligos or HiFi Assembly of single-stranded oligos, respectively. pBbS2k-RFP was a gift from Jay Keasling (Addgene plasmid # 35330 ; http://n2t.net/addgene:35330 ; RRID:Addgene_35330) To clone pSECRETS-B derivatives containing off-target sequences and standard gRNAs or x-gRNA libraries), single-stranded oligonucleotides (oPools) were synthesized by IDT containing the off-target sequences, pLTetO-1 promoter, (N8 random nucleotides immediately upstream) the spacer sequence. These oligos were of the form:

~~~
5’-
CCACTGCTTACTGGCTTATCGGAAGGGATCGTCCTGACCCCG
[Off-target sequence, 20 nt + 3 nt PAM]
CCCCCTCCGTGGAGAAAATTTCCCTATCAGTGATAGAGATTGACATCCCTATCAGTGATAGAGATACTGAGCAC
[5’-extension library, for example: NNNNNNNN]
[20 nt spacer sequence] GTTTTAGAGCTAGAAATAGCAAG
-3’.
~~~

These inserts were PCR’ed with primers

SECRETS-FwdUSER `5’-GCAAG\deoxyU\TAAAATAAGGCTAGTCCG-3’and`

SECRETS-RevUSER `5’-CTTGC\deoxyU\ATTTCTAGCTCTAAAAC-3’`

where deoxyU is a deoxyuracil modified, and the plasmid pSECRETS-B PCR’ed with primers:

Bv3-FwdUSER `5’-GCAAG\deoxyU\TAAAATAAGGCTAGTCCG-3’ and`

B-RevUSER `5’-GTGGG\deoxyU\TCTCTAGTTAGCCAGAG-3’`

These inserts were then cloned into the pSECRETS-B cassette *via* USER cloning (NEB #M5505S). To maintain library diversity, after transformation, E. coli was recovered in 1 mL SOC media for 1 hour without selection, then 0.5 mL of the media was reinoculated into 7 mL LB with kanamycin (50 μg/mL) and grown overnight. 5 mL of the transformants were then centrifuged, then miniprepped.

The pSECRETS-C plasmids containing desired on-target sequences were constructed *via* HiFi assembly into the p11.LacY.wtx1 plasmid (Addgene #69056), which was double digested with XbaI and SphI, with a dsDNA fragment containing a target site (20bp + PAM), 15 bp genomic context sequences flanking the target, and overhang sequences matching the digested plasmid. These fragments were constructed by PCRing short oligos with the form:

~~~
5’-ATAACAGGGTAATATCACGC
[15 bp upstream genomic sequence context] +
[20 bp target sequence + 3 bp PAM] +
[15 bp downstream genomic sequence context]
AAGCTTGGCTGTTTTGGCGG -3’
~~~

*E. coli* strains containing pSECRETS-C plasmids were grown in solutions supplemented with glucose (glu) to suppress leakage of arabinose-induced promoter until selection.

### The SECRETS protocol and analysis

#### Validating selection strength using standard gRNAs

For validation, pSECRETS-C itself or pSECRETS-C containing the *EMX1* target site and flanking sequences (pSECRETS-C-EMX1); pSECRETS-A; and pSECRETS-B to express the standard *EMX1* gRNA (pSECRETS-B-EMX1-gRNA) or pSECRETS-B-EMX1-gRNA containing with *EMX1* OT1 were electroporated sequentially into electrocompetent NEB10beta *E. coli* cells and recovered in SOC media. For the last transformation with pSECRETS-B, recovery media was supplemented with 10 ng/mL aTc for pre-induction of sgRNA and Cas9. Following recovery for 1 hr, cells were plated on LB agar plates under selective (aTc, arabinose, chloramphenicol, kanamycin) and non-selective (glucose, chloramphenicol, kanamycin, ampicillin) conditions and incubated for 24 hours.

### Selection of extended g-RNAs (SECRETS protocol)

pSECRETS-B plasmids containing x-gRNA libraries and the off-target site were screened similarly to the validation experiments with few changes. *E. coli* cells were transformed in two steps instead of three:

*E. coli* strains containing pSECRETS-A plasmids were electroporated with corresponding B and C plasmid simultaneously (75 ng each DNA). Following recovery, cells were centrifuged at 4°C and supernatant was replaced with fresh LB before inoculating 0.5 mL of the culture into 7 mL liquid LB for selective or non-selective conditions and grown overnight. After miniprep (NEB #T1010L) of the resulting cultures, samples were PCR’ed across the gRNA segment and prepared for Illumina next-generation sequencing.

### Analysis of SECRETS outcomes

Small-scale (at least 50,000 reads) next-generation (Amplicon-EZ; Azenta Inc.) was performed of samples from the SECRETS assay. Custom code was written in MATLAB (Mathworks; Natick, MA) to extract and count the 5’-extensions from the x-gRNA sequence of each read; however, in principle, a short line of code can be written to the same effect following the approach found in Reference ^20^. The number of unique 5’-extensions were enumerated per sample and normalized to the total number of reads per sample and averaged across technical replicates (n = 2). The normalized number of reads per 5’-extension were then averaged across biological replicates (n = 2), sorted from most prevalent to least, and the top five most prevalent 5’-extensions per gRNA selected for further characterization.

### *In vitro* validation of x-gRNAs

#### Cas9 ribonucleoprotein (RNP) generation

DNA oligos of sgRNAs and x-gRNAs were designed according to the EnGen sgRNA Synthesis Kit (NEB #E3322) to add 5’-T7 RNA polymerase promoter sequence and 3’-Cas9 crRNA sequence and were purchased from Integrated DNA Technologies IDT then resuspended to a stock concentration of 100 μM. If the (x-)gRNA did not have an initial 5’-dG necessary for T7 RNA polymerase transcription, one was added in the DNA oligo sequence. For sgRNA synthesis, oligos were diluted 100x (1 μM) then used with the EnGen sgRNA Synthesis Kit per manufacturer’s instructions. Cas9 RNPs were formed following the IDT Alt-R CRISPR-Cas9 System – *In vitro* cleavage of target DNA with ribonucleoprotein complex protocol (Option 2). Cas9 enzyme (Sigma Aldrich, #CAS9PROT-250UG), eCas9 enzyme (Sigma Aldrich #ESPCAS9PRO-50UG), or dCas9 enzyme (IDT Alt-R® S.p. dCas9 Protein V3 #1081066) and sgRNA were combined in equimolar amounts in Phosphate buffered saline, pH 7.4 - PBS (ThermoFisher, #10010023) and incubated at room temperature for 10 minutes. Following incubation, RNPs were stored at -80°C or immediately used for *in vitro* digestion reactions.

### *In vitro* digestion reactions

Three hundred (300) bp DNA targets containing the target sequence ∼200 bp from one end and the flanking genomic context were synthesized by Twist Bioscience, PCR amplified using the provided universal primers, purified, and resuspended in nuclease-free water to 100 nM. Three technical replications of reactions were assembled in the following order: 7 μL nuclease-free water, 1 μL target DNA substrate (100 nM), 1 μL 10x Cas9 Nuclease Reaction buffer (200 mM HEPES, 1 M NaCl, 50 mM MgCl2, 1 mM EDTA (pH 6.5 at 25°C)), 1 μL Cas9-RNP (1 mM), then incubated for 1 hour at 37°C followed by proteinase K digestion (1 μL - 56°C for 10 minutes; ThermoFisher, #EO0491). Products were resolved on a 3% agarose gel stained with SYBR Gold and analyzed using ImageJ.

### Evaluation of gene activity of transfected Cas9/x-gRNAs into human cell lines

Cells were transfected using the Lipofectamine CRISPRMAX Cas9 Transfection Reagent (ThermoFisher #CMAX00003) kit. Prior to transfection, A549 fibroblast cells were plated in 24-well plates at 25% confluency in Dulbecco’s Modified Eagle’s Medium - DMEM (ATCC #30-2002) + 10% Fetal Bovine Serum - FBS (ATCC 30-2020) + 1% Penicillin-Streptomycin solution (ATCC 30-2300) and incubated for 24 hours at 37°C + 5% CO_2_. Following incubation, the media was removed and cells were washed with 1x PBS and replaced with fresh DMEM + 10% FBS. Cas9 RNP complexes were formed in a 1:1.2 molar ratio of Cas9 protein to sgRNA with Cas9 Plus reagent to a total volume of 25 μL in Opti-MEM Reduced Serum Medium (ThermoFisher #31985070) per reaction (n = 3). RNPs were added to a mix of 25 μL Opti-MEM I and 1.5 μL CRISPRMAX reagent per reaction, and following a 10 minute room temperature incubation, 50 μL was added to each well. Cells were then incubated at 37°C + 5% CO_2_ for 48 hours.

### Analysis of gene editing outcomes

Cells were processed as follows using the GeneArt Genomic Cleavage Detection Kit (ThermoFisher #A24372). Cell media was collected in a 1.5 mL Eppendorf tube. Remaining attached cells were washed with PBS then detached using TrypLe Express (ATCC 30-2300) trypsin and transferred to the corresponding Eppendorf tube for centrifugation at 1,200 x g at 4°C to pellet cells. Once the supernatant was discarded, pellets were resuspended in cell lysis buffer with protein degrader (supplied in kit) and incubated at 68°C then 95°C to lyse cells. Crude cell lysate was mixed with forward and reverse primer (10 μM), AmpliTaq Gold 360 master mix, and nuclease-free water for direct PCR amplification of the region of interest followed by agarose gel electrophoresis to confirm expected PCR length. Heteroduplexes of the PCR products were formed by mixing with 10x detection buffer and heating samples to 95°C, cooling to 85°C at 2°C/sec, then to 25°C at 0.1°C/sec. Detection enzyme was added to the samples and incubated at 37°C, then fragments were resolved on a 3% agarose gel stained with SYBR Gold. Fluorescence was measured through ImageJ, intensity normalized by length of the DNA fragments, and fraction cleaved was determined using the following equation:

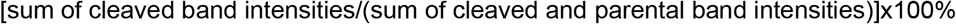

Two technical replicates of samples were also prepared for illumina next-generation sequencing and amplicon sequencing with editing efficiency determined using the CRISPResso2 pipeline.^21^

### Genome-wide off-target screens

We measured genome-wide off-target editing using CHANGE-seq described by Lazzarotto *et al*.^5^ with minor modifications. The genomic DNA purification steps were carried out using the NEB Monarch Genomic DNA Purification Kit. An agarose gel was used to visualize the tagmentation of the human genomic DNA with the transposase. For the PCR step after cleavage and USER enzyme treatment (step 25 of the supplemental information), NEB 2X Q5 Master Mix was used in place of 2X Kapa HiFi HotStart Ready Mix. In place of the MiSeq protocol described in the supplemental information, cleaved genomic DNA barcoded and amplified *via* PCR for illumina sequencing was sent for NGS Amplicon-EZ sequencing by Azenta.

## Availability of code

Short MATLAB scripts used to extract 5’-extension sequences from NGS data will be provided upon request to the corresponding author.

## Availability of materials

pSECRETS-A and pSECRETS-B precursors (without gRNA spacers or off-targets) will be provided to Addgene plasmid repository or made available upon reasonable request to the corresponding author. pSECRETS-C precursor plasmids (without on-target sequences) are available from Addgene (Addgene plasmid # 69056 ; http://n2t.net/addgene:69056 ; RRID:Addgene_69056) as a gift from Huimin Zhao.

## Acknowledgements

We thank Dr. Rachel Tinker-Kulberg and Dr. Cicera R. Lazzarotto for expert technical advice, and members of the Josephs laboratory for careful reading of the manuscript. The work was funded by grants from the National Institute of General Medical Sciences (R35GM133483) and the National Institute of Bioengineering and Biomedical Imaging (R21EB033595) of the National Institutes of Health to EAJ. This work was performed in part at the Joint School of Nanoscience and Nanoengineering, a member of the Southeastern Nanotechnology Infrastructure Corridor (SENIC) and National Nanotechnology Coordinated Infrastructure (NNCI), which is supported by the National Science Foundation (Grant ECCS-1542174).

## Declaration of interests

AHN, HD, MRR, and EAJ are inventors filed by the University of North Carolina at Greensboro for related technologies. EAJ is an inventor in on a patent related to hp-gRNAs and other patent applications related to CRISPR technologies.

