## Supplementary Information for "Selection of Extended CRISPR RNAs with Enhanced Targeting and Specificity"

**SUPPORTING INFORMATION**

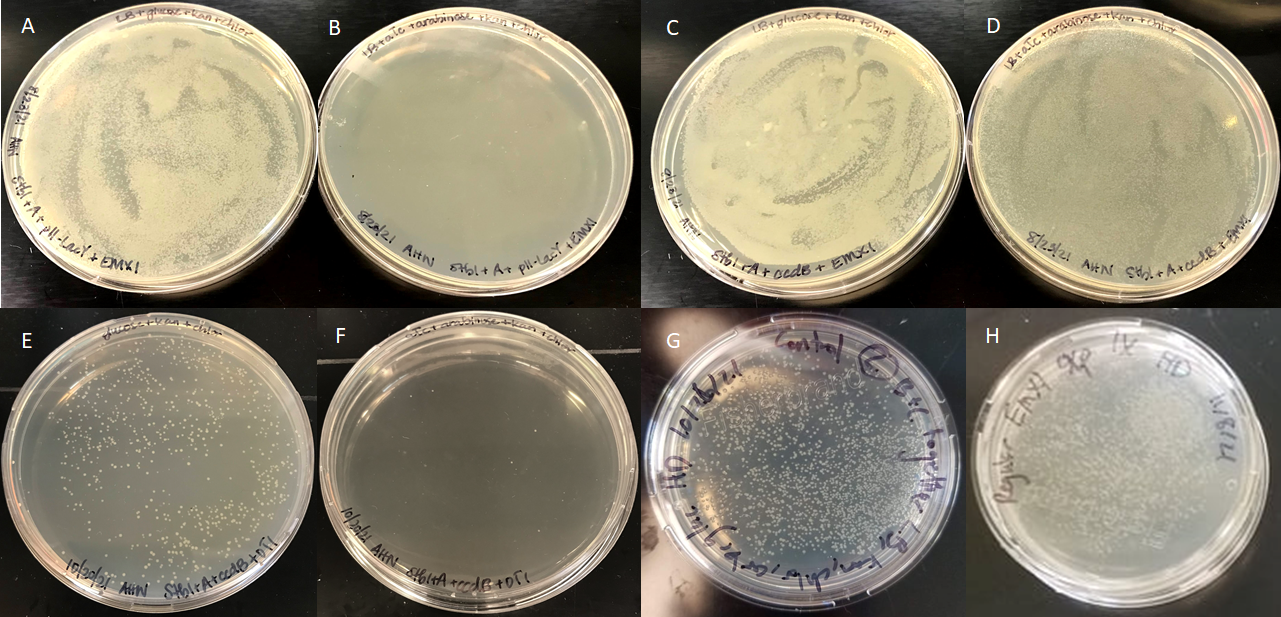
**Figure S1. SECRETS is highly selective for on-target activity and counter-selective for off-target activity.** A and B) SECRETS screen with pSECRETS-A, an empty toxin plasmid (pSECRETS-C with no target), and pSECRETS-B with an *EMX1* sgRNA after plating on non-selective (A) vs selective (B) conditions. C and D) SECRETS screen with pSECRETS-A, pSECRETS-C with *EMX1* ON, and pSECRETS-B with an *EMX1* sgRNA on non-selective (C) vs selective (D) conditions. E and F) SECRETS screen with pSECRETS-A, pSECRETS-C with *EMX1* ON, and pSECRETS-B with an *EMX1* sgRNA and *EMX1* OFF1 on non-selective (E) vs selective (F) conditions. G and H) SECRETS screen with pSECRETS-A, pSECRETS-C with *EMX1* ON, and pSECRETS-B with an *EMX1* x-gRNA library (N8) and *EMX1* OFF1 on non-selective (G) vs selective (H) conditions.

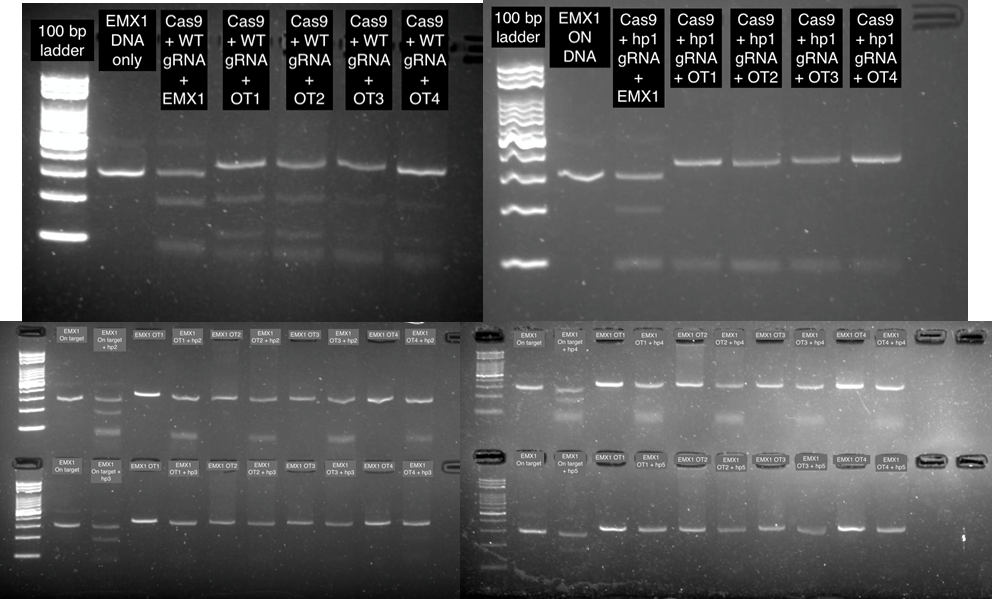

**Figure S2. Representative gels of Cas9 RNP cleavage activity with EMX1 sgRNA and x-gRNAs.**

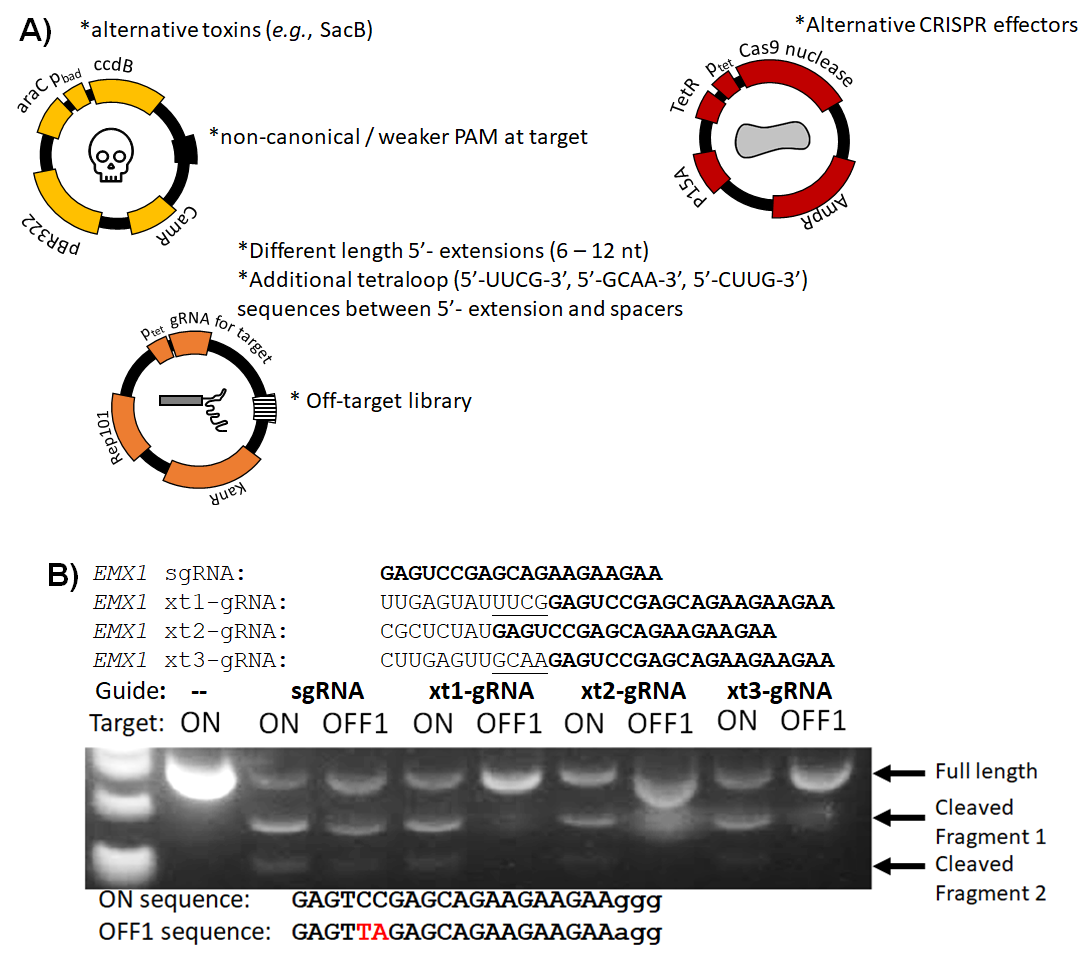

**Figure S3. Modifications to the SECRETS plasmids to expand the 5’-extension space and selection strength.** A) Examples of potential modifications to the SECRETS plasmids. B) *EMX1* x-gRNA libraries with 8 random nucleotides and 4 potential (tetraloop) sequences between N8 and the spacer (UUCG, GCAA, CUUG, or no tetraloop sequence) were screened against *EMX1* OFF1 using SECRETS assay—4^8 * 4 = 262,144 possible 5’- extensions. 3 colonies of the survivors were picked at random and sequenced using colony PCR and Sanger sequencing then characterized using in vitro cleavage assays of purified RNPs, and all 3 exhibited extremely strong on-target activity (comparable to sgRNA) and essentially no off-target activity at *EMX1* OFF1.

**Table S1: Top 50 5’- extensions from SECRETS protocol for EMX1, FANCF, and VEGFA targets and top off-target.**

| ***EMX1*** | | ***VEGFA*** | | ***FANCF*** | |
| --- | --- | --- | --- | --- | --- |
| **5'- extension sequences** | **Fraction reads** | **5'- extension sequences** | **Fraction reads** | **5'- extension sequences** | **Fraction reads** |
| ATGCTTCA | 0.063642 | GGCCCAGC | 0.022094 | ACATTCCT | 0.028027 |
| AAGTACGG | 0.044337 | ATCGCGAT | 0.015392 | AAAGATAG | 0.01289 |
| AAGAGTTA | 0.033594 | TTTGGAAT | 0.013769 | AATTACTT | 0.007923 |
| AGGGATAG | 0.024857 | TCTCACCA | 0.011858 | CCAAGTCT | 0.007864 |
| TGGCTATG | 0.020792 | GGGCCCTC | 0.0089 | ATTGGTGA | 0.0068 |
| CAATAAAG | 0.018731 | GGTGTGGC | 0.008508 | GTCGTTAC | 0.005144 |
| AAAGTCTG | 0.018671 | CCTTTGAC | 0.008036 | ACGACGTA | 0.004849 |
| GAGATCAG | 0.016166 | CTACTCCA | 0.007487 | GAAGATTG | 0.004789 |
| ATGGCAGG | 0.011847 | TAACTATC | 0.007356 | CAATAAAA | 0.004435 |
| GTTAGTAG | 0.011538 | CCGCCGAA | 0.00733 | GAGTGCTC | 0.004316 |
| ACGAGTGT | 0.011184 | TGCGTAGC | 0.006937 | GGGTGAAC | 0.004316 |
| TGCGTGAG | 0.009982 | TGAACTAA | 0.006701 | ATGGCCGG | 0.004198 |
| CGTGGTAT | 0.009542 | TATGTCCC | 0.006335 | GCCCACAA | 0.00408 |
| GATTAGCG | 0.0095 | TCGACTTC | 0.006021 | ATAGGTGT | 0.003548 |
| AGTGGATC | 0.009345 | CCGCATCC | 0.005942 | GGCCATAC | 0.003548 |
| AAACTAGC | 0.008774 | TGGGGCGC | 0.005942 | GTGCAATT | 0.003548 |
| GGTGAGAG | 0.008427 | CCTCCTGC | 0.005864 | GACGAGCC | 0.00343 |
| TCTCGGTG | 0.007407 | TTCATTTA | 0.005811 | GAGGCGTT | 0.003311 |
| TAAATTTT | 0.00716 | CCCTTTCA | 0.005759 | GAGGTAAC | 0.003311 |
| TGGTCCTC | 0.007007 | TGCCTGCA | 0.005733 | GTCACGCT | 0.003311 |
| GGTATTTT | 0.006975 | ATGCATAG | 0.005654 | ATTCGAGT | 0.003252 |
| GGTGCATA | 0.006914 | CCAGCGAA | 0.005602 | GGCGTCTT | 0.003193 |
| GAGAGCAA | 0.006854 | GGTGAGGC | 0.005602 | GGGCACAG | 0.003193 |
| TGACCCAC | 0.006641 | TGCCCACT | 0.005366 | GGATGAGG | 0.003016 |
| GGCGCCGT | 0.006529 | CCCTTGTA | 0.005026 | CCTCTGGT | 0.002956 |
| CTGCTCCT | 0.006518 | TAATCTCT | 0.005 | GCAACCCT | 0.002956 |
| AATCAAGC | 0.006449 | CCGGCATT | 0.004895 | ATTGGTCT | 0.002838 |
| AATCACCG | 0.0064 | GATGTAGG | 0.004843 | AATAAAAC | 0.00272 |
| TTATCAGT | 0.006298 | GTACTATC | 0.004712 | ACCTTTAA | 0.00272 |
| AGATTTTC | 0.006248 | ATGCGATA | 0.00466 | GCCTGGAC | 0.002602 |
| TTTGTAGA | 0.006229 | TTCTACAC | 0.004633 | TAGTCATC | 0.002602 |
| GTATGGTG | 0.006108 | CCAGCCAT | 0.004581 | AGATGCAT | 0.002543 |
| GGCAGGTG | 0.006082 | GCTTCGCC | 0.004424 | AGGCGAGA | 0.002543 |
| GGTAAGTT | 0.006025 | CCCTGCCG | 0.004372 | ATGTTCGA | 0.002543 |
| GTGCACGT | 0.005758 | ACCCTGGC | 0.004293 | CAGATTTC | 0.002483 |
| ACAATTGA | 0.005685 | GGGCAATG | 0.004293 | CCCTATGT | 0.002483 |
| AGGTGTGA | 0.005567 | TGAGGATG | 0.004267 | GCGTAGTG | 0.002483 |
| GTCGTGCC | 0.005317 | CAGGCTCA | 0.004241 | GGTATTTT | 0.002365 |
| ATTAGAGT | 0.005264 | GGGTAGGA | 0.004215 | AGATTTAA | 0.002306 |
| CCGCCTTT | 0.005216 | CCCCCGTC | 0.004057 | GATCCGTT | 0.002247 |
| TGTGTTTC | 0.005119 | TCAGAATG | 0.003979 | GGCTAATC | 0.002247 |
| CACTGGTC | 0.004938 | CCAGCAAC | 0.003953 | GTGCTTTG | 0.002247 |
| TGACCAAG | 0.004926 | ACTAAAAA | 0.003927 | ATAGTTTT | 0.002188 |
| TAACGATG | 0.00489 | AAGCGGGA | 0.003796 | ATCATTGA | 0.002129 |
| TGACGGGT | 0.00477 | ACTAGCTC | 0.003796 | GCGCTACG | 0.002129 |
| GAGGTAGG | 0.004761 | GCGTGTGA | 0.003796 | ACTGGTGT | 0.001951 |
| GGGTTTAC | 0.004753 | GGGGTGGC | 0.00377 | ATATTGGG | 0.001951 |
| GACGTTAG | 0.004693 | CCCACTGT | 0.003691 | GACTCTAA | 0.001951 |
| TCTTTACG | 0.004604 | GTCCCCAC | 0.003691 | ATCATTTC | 0.001892 |

**Table S2. Oligonucleotide and primer sequences**

| **Primer Name** | **Sequence** | **Assay** |
| --- | --- | --- |
| pBbA2K-RFP-Fwd | AGTCACACTGGCTCACCTTC | Cloning (pSECRETS-A) |
| pBbA2K-RFP-Rev | TAACAACCCGTAAACTCGCC | Cloning (pSECRETS-A) |
| Cas9-Fwd | GCACAAATAGCGTCGGATGG | Cloning (pSECRETS-A) |
| Cas9-Rev | GAAGGTGAGCCAGTGTGACT | Cloning (pSECRETS-A) |
| gRNA-3 | TCGTTGTGGGAGGTGATGTC | Cloning (pSECRETS-A) |
| gRNA-5 | GACGAAGACTCAATTG | Cloning (pSECRETS-B) |
| pBbS2k/gRNA-fwd | GACATCACCTCCCACAACGA | Cloning (pSECRETS-B) |
| pBbS2k/gRNA-rev | GGTAAGGTTGGGAAGCCCTG | Cloning (pSECRETS-B) |
| pBbS2k/gRNA-check5 | CCTGCGCCATCAGATCCTT | Cloning (pSECRETS-B) |
| pBbS2k/gRNA-check3 | GGTGGAGTGACGACCTTCAG | Cloning (pSECRETS-B) |
| pBbS2k-fwd | GACATCACCTCCCACAACGA | Cloning (pSECRETS-B) |
| pBbS2k-rev | TAACAACCCGTAAACTCGCC | Cloning (pSECRETS-B) |
| p11-Fwd | AAG CTT GGC TGT TTT GGC G | Cloning (pSECRETS-C) |
| p11-Rev | GCG TGA TAT TAC CCT GTT ATC CC | Cloning (pSECRETS-C) |
| Target-Fwd | ATA ACA GGG TAA TAT CAC GC | Cloning (pSECRETS-C) |
| EMX1-Rev | CCG CCA AAA CAG CCA AGC TTT TGA TGT GAT GGG AGC TCT T | Cloning (pSECRETS-C) |
| FANCF-Rev | CCG CCA AAA CAG CCA AGC TTG CTC GGA AAA GCG ATC TAG | Cloning (pSECRETS-C) |
| VEGFA-Rev | CCG CCA AAA CAG CCA AGC TTT CAG AAA TAG GGG GTC CAG | Cloning (pSECRETS-C) |
| B-FwdUSER | GCAAG\deoxyU\TAAAATAAGGCTAGTCCG | Cloning (x-)gRNA libraries and off-targets into pSECRETS-B |
| B-RevUSER | GTGGG\deoxyU\TCTCTAGTTAGCCAGAG | Cloning (x-)gRNA libraries and off-targets into pSECRETS-B |
| SECRETS-FwdUSER | CCCAC\deoxyU\GCTTACTGGCTTATCG | Cloning (x-)gRNA libraries and off-targets into pSECRETS-B |
| SECRETS-RevUSER | CTTGC\deoxyU\ATTTCTAGCTCTAAAAC | Cloning (x-)gRNA libraries and off-targets into pSECRETS-B |
| EMX1-OFF1-N8-xgRNA | CCACTGCTTACTGGCTTATCGGAAGGGCAAGACAGATTGTCAGAGTTAGAGCAGAAGAAGAAAGGCATGGAGTAAAGGCAAAATTTCCCTATCAGTGATAGAGATTGACATCCCTATCAGTGATAGAGATACTGAGCACNNNNNNNNGAGTCCGAGCAGAAGAAGAAGTTTTAGAGCTAGAAATAGCAAGTTAAA | Cloning EMX1 OFF1 and N8 x-gRNA library into pSECRETS-B |
| EMX1-OFF1(noPAM)-N8-xgRNA | CCACTGCTTACTGGCTTATCGGAAGGGCAAGACAGATTGTCAGAGTTAGAGCAGAAGAAGAATTTCATGGAGTAAAGGCAAAATTTCCCTATCAGTGATAGAGATTGACATCCCTATCAGTGATAGAGATACTGAGCACNNNNNNNNGAGTCCGAGCAGAAGAAGAAGTTTTAGAGCTAGAAATAGCAAGTTAAA | Cloning EMX1 (no OFF1) N8 x-gRNA library into pSECRETS-B |
| FANCF-OFF1-N8-xgRNA | CCACTGCTTACTGGCTTATCGgaagggatcgtcctgaccccgggaaccccgtctgcagcaccAGGccccctccgtggagaAaatttccctatcaGTGATAGagattgacatccctatcagtgatagagatactgagcacNNNNNNNNggaatcccttctgcagcaccgttttagagctagaaatagcaag | Cloning FANCF OFF1 and N8 x-gRNA library into pSECRETS-B |
| FANCF-OFF1(noPAM)-N8-xgRNA | CCACTGCTTACTGGCTTATCGgaagggcttcgcgcacctcatggaatcccttctgcagcaccTTTatcgcttttccgagcAaatttccctatcagtgatagagattgacatccctatcagtgatagagatactgagcacNNNNNNNNggaatcccttctgcagcaccgttttagagctagaaatagcaag | Cloning FANCF (no OFF1) N8 x-gRNA library into pSECRETS-B |
| VEGFA-OFF1-N8-xgRNA | CCACTGCTTACTGGCTTATCGgaagggaaggacggatttgtgggatggagggagtttgctccTGGggtgtcagaatgtccAaatttccctatcagtgatagagattgacatccctatcagtgatagagatactgagcacNNNNNNNNgggtggggggagtttgctccgttttagagctagaaatagcaag | Cloning VEGFA OFF1 and N8 x-gRNA library into pSECRETS-B |
| VEGFA-OFF1(noPAM)-N8-xgRNA | CCACTGCTTACTGGCTTATCGgaagggtgggctttggaaagggggtggggggagtttgctccTTTaccccctatttctgaAaatttccctatcagtgatagagattgacatccctatcagtgatagagatactgagcacNNNNNNNNgggtggggggagtttgctccgttttagagctagaaatagcaag | Cloning VEGFA (no OFF1) N8 x-gRNA library into pSECRETS-B |
| EMX1-Fwd | TATGTAGCCTCAGTCTTCCCATCA | T7E1 mutation detection |
| EMX1-Rev | TGGTTGCCCACCCTAGTCATTG | T7E1 mutation detection |
| EMX1-OT1-Fwd | GAAAACTCAAAGAAATGCCCAATCA | T7E1 mutation detection |
| EMX1-OT1-Rev | TCTGTGGAGGCAATAAAGTGAAATAA | T7E1 mutation detection |
| FANCF-Fwd | GGGCCTGGAAGTTCGCTAAT | T7E1 mutation detection |
| FANCF-Rev | AGGCGTATCATTTCGCGGAT | T7E1 mutation detection |
| FANCF-OT1-Fwd | GTTGCGTTTGAGGATGAGCA | T7E1 mutation detection |
| FANCF-OT1-Rev | GAAATCTCCCACGAGTCTCCG | T7E1 mutation detection |
| VEGFA-Fwd | TCCAGATGGCACATTGTCAG | T7E1 mutation detection |
| VEGFA-Rev | AGGGAGCAGGAAAGTGAGGT | T7E1 mutation detection |
| VEGFA-OT1-Fwd | ACCCCACAGCCAGGTTTTCA | T7E1 mutation detection |
| VEGFA-OT1-Rev | GAATCACTGCACCTGGCCATC | T7E1 mutation detection |
| EMX1 On fwd 1 | ACACTCTTTCCCTACACGACGCTCTTCCGATCTATCACGAGTTTCTCATCTGTGCCCCT | Illumina Sequencing |
| EMX1 On fwd 2 | ACACTCTTTCCCTACACGACGCTCTTCCGATCTCGATGTAAGTTTCTCATCTGTGCCCCT | Illumina Sequencing |
| EMX1 On fwd 3 | ACACTCTTTCCCTACACGACGCTCTTCCGATCTTTAGGCTAAGTTTCTCATCTGTGCCCCT | Illumina Sequencing |
| EMX1 On fwd 4 | ACACTCTTTCCCTACACGACGCTCTTCCGATCTTGACCAGTAAGTTTCTCATCTGTGCCCCT | Illumina Sequencing |
| EMX1 On fwd 5 | ACACTCTTTCCCTACACGACGCTCTTCCGATCT ACATGT CGTA AGTTTCTCATCTGTGCCCCT | Illumina Sequencing |
| EMX1 On fwd 6 | ACACTCTTTCCCTACACGACGCTCTTCCGATCT GCCAAT ACGTA AGTTTCTCATCTGTGCCCCT | Illumina Sequencing |
| EMX1 On rev 1 | GACTGGAGTTCAGACGTGTGCTCTTCCGATCT CAGATC ATCGATGTCCTCCCCATTGG | Illumina Sequencing |
| EMX1 On rev 2 | GACTGGAGTTCAGACGTGTGCTCTTCCGATCT ACTTGA ATCGATGTCCTCCCCATTGG | Illumina Sequencing |
| EMX1 On rev 3 | GACTGGAGTTCAGACGTGTGCTCTTCCGATCT GATCAG ATCGATGTCCTCCCCATTGG | Illumina Sequencing |
| EMX1 On rev 4 | GACTGGAGTTCAGACGTGTGCTCTTCCGATCT TAGCTT ATCGATGTCCTCCCCATTGG | Illumina Sequencing |
| EMX1 On rev 5 | GACTGGAGTTCAGACGTGTGCTCTTCCGATCT GGCTAG ATCGATGTCCTCCCCATTGG | Illumina Sequencing |
| EMX1 On rev 6 | GACTGGAGTTCAGACGTGTGCTCTTCCGATCT CTTGTA ATCGATGTCCTCCCCATTGG | Illumina Sequencing |
| EMX1 OT1 fwd 1 | ACACTCTTTCCCTACACGACGCTCTTCCGATCT ATCACG CGCTTGTCCATGTCTAGGAA | Illumina Sequencing |
| EMX1 OT1 fwd 2 | ACACTCTTTCCCTACACGACGCTCTTCCGATCT CGATGT A CGCTTGTCCATGTCTAGGAA | Illumina Sequencing |
| EMX1 OT1 fwd 3 | ACACTCTTTCCCTACACGACGCTCTTCCGATCT TTAGGC TA CGCTTGTCCATGTCTAGGAA | Illumina Sequencing |
| EMX1 OT1 fwd 4 | ACACTCTTTCCCTACACGACGCTCTTCCGATCT TGACCA GTA CGCTTGTCCATGTCTAGGAA | Illumina Sequencing |
| EMX1 OT1 fwd 5 | ACACTCTTTCCCTACACGACGCTCTTCCGATCT ACATGT CGTA CGCTTGTCCATGTCTAGGAA | Illumina Sequencing |
| EMX1 OT1 fwd 6 | ACACTCTTTCCCTACACGACGCTCTTCCGATCT GCCAAT ACGTA CGCTTGTCCATGTCTAGGAA | Illumina Sequencing |
| EMX1 OT1 rev 1 | GACTGGAGTTCAGACGTGTGCTCTTCCGATCT CAGATC TGGCATGGCAAGACAGATTG | Illumina Sequencing |
| EMX1 OT1 rev 2 | GACTGGAGTTCAGACGTGTGCTCTTCCGATCT ACTTGA TGGCATGGCAAGACAGATTG | Illumina Sequencing |
| EMX1 OT1 rev 3 | GACTGGAGTTCAGACGTGTGCTCTTCCGATCT GATCAG TGGCATGGCAAGACAGATTG | Illumina Sequencing |
| EMX1 OT1 rev 4 | GACTGGAGTTCAGACGTGTGCTCTTCCGATCT TAGCTT TGGCATGGCAAGACAGATTG | Illumina Sequencing |
| EMX1 OT1 rev 5 | GACTGGAGTTCAGACGTGTGCTCTTCCGATCT GGCTAG TGGCATGGCAAGACAGATTG | Illumina Sequencing |
| EMX1 OT1 rev 6 | GACTGGAGTTCAGACGTGTGCTCTTCCGATCT CTTGTA TGGCATGGCAAGACAGATTG | Illumina Sequencing |
| FANCF On fwd 1 | ACACTCTTTCCCTACACGACGCTCTTCCGATCT ATCACG CTCCAGAGCCGTGCGAATG | Illumina Sequencing |
| FANCF On fwd 2 | ACACTCTTTCCCTACACGACGCTCTTCCGATCT CGATGT A CTCCAGAGCCGTGCGAATG | Illumina Sequencing |
| FANCF On fwd 3 | ACACTCTTTCCCTACACGACGCTCTTCCGATCT TTAGGC TA CTCCAGAGCCGTGCGAATG | Illumina Sequencing |
| FANCF On fwd 4 | ACACTCTTTCCCTACACGACGCTCTTCCGATCT TGACCA GTA CTCCAGAGCCGTGCGAATG | Illumina Sequencing |
| FANCF On fwd 5 | ACACTCTTTCCCTACACGACGCTCTTCCGATCT ACATGT CGTA CTCCAGAGCCGTGCGAATG | Illumina Sequencing |
| FANCF On fwd 6 | ACACTCTTTCCCTACACGACGCTCTTCCGATCT GCCAAT ACGTA CTCCAGAGCCGTGCGAATG | Illumina Sequencing |
| FANCF On rev 1 | GACTGGAGTTCAGACGTGTGCTCTTCCGATCT CAGATC AATCAGTACGCAGAGAGTCGC | Illumina Sequencing |
| FANCF On rev 2 | GACTGGAGTTCAGACGTGTGCTCTTCCGATCT ACTTGA AATCAGTACGCAGAGAGTCGC | Illumina Sequencing |
| FANCF On rev 3 | GACTGGAGTTCAGACGTGTGCTCTTCCGATCT GATCAG AATCAGTACGCAGAGAGTCGC | Illumina Sequencing |
| FANCF On rev 4 | GACTGGAGTTCAGACGTGTGCTCTTCCGATCT TAGCTT AATCAGTACGCAGAGAGTCGC | Illumina Sequencing |
| FANCF On rev 5 | GACTGGAGTTCAGACGTGTGCTCTTCCGATCT GGCTAG AATCAGTACGCAGAGAGTCGC | Illumina Sequencing |
| FANCF On rev 6 | GACTGGAGTTCAGACGTGTGCTCTTCCGATCT CTTGTA AATCAGTACGCAGAGAGTCGC | Illumina Sequencing |
| FANCF OT1 fwd 1 | ACACTCTTTCCCTACACGACGCTCTTCCGATCT ATCACG AGTTGGTGTCAATACAACTCCCCT | Illumina Sequencing |
| FANCF OT1 fwd 2 | ACACTCTTTCCCTACACGACGCTCTTCCGATCT CGATGT A AGTTGGTGTCAATACAACTCCCCT | Illumina Sequencing |
| FANCF OT1 fwd 3 | ACACTCTTTCCCTACACGACGCTCTTCCGATCT TTAGGC TA AGTTGGTGTCAATACAACTCCCCT | Illumina Sequencing |
| FANCF OT1 fwd 4 | ACACTCTTTCCCTACACGACGCTCTTCCGATCT TGACCA GTA AGTTGGTGTCAATACAACTCCCCT | Illumina Sequencing |
| FANCF OT1 fwd 5 | ACACTCTTTCCCTACACGACGCTCTTCCGATCT ACATGT CGTA AGTTGGTGTCAATACAACTCCCCT | Illumina Sequencing |
| FANCF OT1 fwd 6 | ACACTCTTTCCCTACACGACGCTCTTCCGATCT GCCAAT ACGTA AGTTGGTGTCAATACAACTCCCCT | Illumina Sequencing |
| FANCF OT1 rev 1 | GACTGGAGTTCAGACGTGTGCTCTTCCGATCT CAGATC CGCCAGCACTTTCTAAGGAATC | Illumina Sequencing |
| FANCF OT1 rev 2 | GACTGGAGTTCAGACGTGTGCTCTTCCGATCT ACTTGA CGCCAGCACTTTCTAAGGAATC | Illumina Sequencing |
| FANCF OT1 rev 3 | GACTGGAGTTCAGACGTGTGCTCTTCCGATCT GATCAG CGCCAGCACTTTCTAAGGAATC | Illumina Sequencing |
| FANCF OT1 rev 4 | GACTGGAGTTCAGACGTGTGCTCTTCCGATCT TAGCTT CGCCAGCACTTTCTAAGGAATC | Illumina Sequencing |
| FANCF OT1 rev 5 | GACTGGAGTTCAGACGTGTGCTCTTCCGATCT GGCTAG CGCCAGCACTTTCTAAGGAATC | Illumina Sequencing |
| FANCF OT1 rev 6 | GACTGGAGTTCAGACGTGTGCTCTTCCGATCT CTTGTA CGCCAGCACTTTCTAAGGAATC | Illumina Sequencing |

**Table S3. dsDNA fragments used to clone pSECRETS plasmids.**

| pSECRETS-A gBlock | GGCGAGTTTACGGGTTGTTAaaccttcgattccgacctcattaagcagctctaatgcgctgttaatcactttacttttatctaaacgagacatactcttcctttttcaatattattgaagcatttatcagggttattgtctcatgagcggatacatatttgaatgtatttagaaaaataaacaaataggggttccgcgcacatttccccgaaaagtgccacctgacgtcctctggctaactagagaacccactgcttactggcttatcgaaatttccctatcagtgatagagattgacatccctatcagtgatagagatactgagcacatcagcaggacgcactgaccagggagacccaagcttgccaccatggtgtacccctacgacgtgcccgactacgccgaattgcctccaaaaaagaagagaaaggtagggatccgaaccatggataagaaatactcaataggcttagatatcgGCACAAATAGCGTCGGATGG | TetR-pAmp-pltetO1-Cas9 for pBbA2c |
| --- | --- | --- |
| pSECRETS-B gBlock | GGCGAGTTTACGGGTTGTTAaaccttcgattccgacctcattaagcagctctaatgcgctgttaatcactttacttttatctaaacgagacatactcttcctttttcaatattattgaagcatttatcagggttattgtctcatgagcggatacatatttgaatgtatttagaaaaataaacaaataggggttccgcgcacatttccccgaaaagtgccacctgacgtcctctggctaactagagaacccactgcttactggcttatcgaaatttccctatcagtgatagagattgacatccctatcagtgatagagatactgagcacgtGAGACCcatgccatagcgttgttTAGGGATAACAGGGTAATaCTGTCCACACAATCTGCCCTGGTCTCcgttttagagctagaaatagcaagttaaaataaggctagtccgttatcaacttgaaaaagtggcaccgagtcggtgcttttttGGCCGGCATGGTCCCAGCCTCCTCGCTGGCGCCGGCTGGGCAACATGCTTCGGCATGGCGAATGGGACGACATCACCTCCCACAACGA | TetR-pAmp-pltetO1-gRNA (BsaI cassette) |

**Table S4. Plasmid Sequences**

**pSECRETS-A:**

gacgtcttaagacccactttcacatttaagttgtttttctaatccgcatatgatcaattcaaggccgaataagaaggctggctctgcaccttggtgatcaaataattcgatagcttgtcgtaataatggcggcatactatcagtagtaggtgtttccctttcttctttagcgacttgatgctcttgatcttccaatacgcaacctaaagtaaaatgccccacagcgctgagtgcatataatgcattctctagtgaaaaaccttgttggcataaaaaggctaattgattttcgagagtttcatactgtttttctgtaggccgtgtacctaaatgtacttttgctccatcgcgatgacttagtaaagcacatctaaaacttttagcgttattacgtaaaaaatcttgccagctttccccttctaaagggcaaaagtgagtatggtgcctatctaacatctcaatggctaaggcgtcgagcaaagcccgcttattttttacatgccaatacaatgtaggctgctctacacctagcttctgGGCGAGTTTACGGGTTGTTAaaccttcgattccgacctcattaagcagctctaatgcgctgttaatcactttacttttatctaaacgagacatactcttcctttttcaatattattgaagcatttatcagggttattgtctcatgagcggatacatatttgaatgtatttagaaaaataaacaaataggggttccgcgcacatttccccgaaaagtgccacctgacgtcctctggctaactagagaacccactgcttactggcttatcgaaatttccctatcagtgatagagattgacatccctatcagtgatagagatactgagcacatcagcaggacgcactgaccagggagacccaagcttgccaccatggtgtacccctacgacgtgcccgactacgccgaattgcctccaaaaaagaagagaaaggtagggatccgaaccatggataagaaatactcaataggcttagatatcgGCACAAATAGCGTCGGATGGgcggtgatcactgatgaatataaggttccgtctaaaaagttcaaggttctgggaaatacagaccgccacagtatcaaaaaaaatcttataggggctcttttatttgacagtggagagacagcggaagcgactcgtctcaaacggacagctcgtagaaggtatacacgtcggaagaatcgtatttgttatctacaggagattttttcaaatgagatggcgaaagtagatgatagtttctttcatcgacttgaagagtcttttttggtggaagaagacaagaagcatgaacgtcatcctatttttggaaatatagtagatgaagttgcttatcatgagaaatatccaactatctatcatctgcgaaaaaaattggtagattctactgataaagcggatttgcgcttaatctatttggccttagcgcatatgattaagtttcgtggtcattttttgattgagggagatttaaatcctgataatagtgatgtggacaaactatttatccagttggtacaaacctacaatcaattatttgaagaaaaccctattaacgcaagtggagtagatgctaaagcgattctttctgcacgattgagtaaatcaagacgattagaaaatctcattgctcagctccccggtgagaagaaaaatggcttatttgggaatctcattgctttgtcattgggtttgacccctaattttaaatcaaattttgatttggcagaagatgctaaattacagctttcaaaagatacttacgatgatgatttagataatttattggcgcaaattggagatcaatatgctgatttgtttttggcagctaagaatttatcagatgctattttactttcagatatcctaagagtaaatactgaaataactaaggctcccctatcagcttcaatgattaaacgctacgatgaacatcatcaagacttgactcttttaaaagctttagttcgacaacaacttccagaaaagtataaagaaatcttttttgatcaatcaaaaaacggatatgcaggttatattgatgggggagctagccaagaagaattttataaatttatcaaaccaattttagaaaaaatggatggtactgaggaattattggtgaaactaaatcgtgaagatttgctgcgcaagcaacggacctttgacaacggctctattccccatcaaattcacttgggtgagctgcatgctattttgagaagacaagaagacttttatccatttttaaaagacaatcgtgagaagattgaaaaaatcttgacttttcgaattccttattatgttggtccattggcgcgtggcaatagtcgttttgcatggatgactcggaagtctgaagaaacaattaccccatggaattttgaagaagttgtcgataaaggtgcttcagctcaatcatttattgaacgcatgacaaactttgataaaaatcttccaaatgaaaaagtactaccaaaacatagtttgctttatgagtattttacggtttataacgaattgacaaaggtcaaatatgttactgaaggaatgcgaaaaccagcatttctttcaggtgaacagaagaaagccattgttgatttactcttcaaaacaaatcgaaaagtaaccgttaagcaattaaaagaagattatttcaaaaaaatagaatgttttgatagtgttgaaatttcaggagttgaagatagatttaatgcttcattaggtacctaccatgatttgctaaaaattattaaagataaagattttttggataatgaagaaaatgaagatatcttagaggatattgttttaacattgaccttatttgaagatagggagatgattgaggaaagacttaaaacatatgctcacctctttgatgataaggtgatgaaacagcttaaacgtcgccgttatactggttggggacgtttgtctcgaaaattgattaatggtattagggataagcaatctggcaaaacaatattagattttttgaaatcagatggttttgccaatcgcaattttatgcagctgatccatgatgatagtttgacatttaaagaagacattcaaaaagcacaagtgtctggacaaggcgatagtttacatgaacatattgcaaatttagctggtagccctgctattaaaaaaggtattttacagactgtaaaagttgttgatgaattggtcaaagtaatggggcggcataagccagaaaatatcgttattgaaatggcacgtgaaaatcagacaactcaaaagggccagaaaaattcgcgagagcgtatgaaacgaatcgaagaaggtatcaaagaattaggaagtcagattcttaaagagcatcctgttgaaaatactcaattgcaaaatgaaaagctctatctctattatctccaaaatggaagagacatgtatgtggaccaagaattagatattaatcgtttaagtgattatgatgtcgatcacattgttccacaaagtttccttaaagacgattcaatagacaataaggtcttaacgcgttctgataaaaatcgtggtaaatcggataacgttccaagtgaagaagtagtcaaaaagatgaaaaactattggagacaacttctaaacgccaagttaatcactcaacgtaagtttgataatttaacgaaagctgaacgtggaggtttgagtgaacttgataaagctggttttatcaaacgccaattggttgaaactcgccaaatcactaagcatgtggcacaaattttggatagtcgcatgaatactaaatacgatgaaaatgataaacttattcgagaggttaaagtgattaccttaaaatctaaattagtttctgacttccgaaaagatttccaattctataaagtacgtgagattaacaattaccatcatgcccatgatgcgtatctaaatgccgtcgttggaactgctttgattaagaaatatccaaaacttgaatcggagtttgtctatggtgattataaagtttatgatgttcgtaaaatgattgctaagtctgagcaagaaataggcaaagcaaccgcaaaatatttcttttactctaatatcatgaacttcttcaaaacagaaattacacttgcaaatggagagattcgcaaacgccctctaatcgaaactaatggggaaactggagaaattgtctgggataaagggcgagattttgccacagtgcgcaaagtattgtccatgccccaagtcaatattgtcaagaaaacagaagtacagacaggcggattctccaaggagtcaattttaccaaaaagaaattcggacaagcttattgctcgtaaaaaagactgggatccaaaaaaatatggtggttttgatagtccaacggtagcttattcagtcctagtggttgctaaggtggaaaaagggaaatcgaagaagttaaaatccgttaaagagttactagggatcacaattatggaaagaagttcctttgaaaaaaatccgattgactttttagaagctaaaggatataaggaagttaaaaaagacttaatcattaaactacctaaatatagtctttttgagttagaaaacggtcgtaaacggatgctggctagtgccggagaattacaaaaaggaaatgagctggctctgccaagcaaatatgtgaattttttatatttagctagtcattatgaaaagttgaagggtagtccagaagataacgaacaaaaacaattgtttgtggagcagcataagcattatttagatgagattattgagcaaatcagtgaattttctaagcgtgttattttagcagatgccaatttagataaagttcttagtgcatataacaaacatagagacaaaccaatacgtgaacaagcagaaaatattattcatttatttacgttgacgaatcttggagctcccgctgcttttaaatattttgatacaacaattgatcgtaaacgatatacgtctacaaaagaagttttagatgccactcttatccatcaatccatcactggtctttatgaaacacgcattgatttgagtcagctaggaggtgactaactcgagtaaggatctccaggcatcaaataaaacgaaaggctcagtcgaaagactgggcctttcgttttatctgttgtttgtcggtgaacgctctctactagAGTCACACTGGCTCACCTTCgggtgggcctttctgcgtttatacctagggatatattccgcttcctcgctcactgactcgctacgctcggtcgttcgactgcggcgagcggaaatggcttacgaacggggcggagatttcctggaagatgccaggaagatacttaacagggaagtgagagggccgcggcaaagccgtttttccataggctccgcccccctgacaagcatcacgaaatctgacgctcaaatcagtggtggcgaaacccgacaggactataaagataccaggcgtttccccctggcggctccctcgtgcgctctcctgttcctgcctttcggtttaccggtgtcattccgctgttatggccgcgtttgtctcattccacgcctgacactcagttccgggtaggcagttcgctccaagctggactgtatgcacgaaccccccgttcagtccgaccgctgcgccttatccggtaactatcgtcttgagtccaacccggaaagacatgcaaaagcaccactggcagcagccactggtaattgatttagaggagttagtcttgaagtcatgcgccggttaaggctaaactgaaaggacaagttttggtgactgcgctcctccaagccagttacctcggttcaaagagttggtagctcagagaaccttcgaaaaaccgccctgcaaggcggttttttcgttttcagagcaagagattacgcgcagaccaaaacgatctcaagaagatcatcttattaatcagataaaatatttctagatttcagtgcaatttatctcttcaaatgtagcacctgaagtcagccccatacgatataagttgttactagtgcttggattctcaccaataaaaaacgcccggcggcaaccgagcgttctgaacaaatccagatggagttctgaggtcattactggatctatcaacaggagtccaagcgagctcgatatcaaattacgccccgccctgccactcatcgcagtactgttgtaattcattaagcattctgccgacatggaagccatcacaaacggcatgatgaacctgaatcgccagcggcatcagcaccttgtcgccttgcgtataatatttgcccatggtgaaaacgggggcgaagaagttgtccatattggccacgtttaaatcaaaactggtgaaactcacccagggattggctgagacgaaaaacatattctcaataaaccctttagggaaataggccaggttttcaccgtaacacgccacatcttgcgaatatatgtgtagaaactgccggaaatcgtcgtggtattcactccagagcgatgaaaacgtttcagtttgctcatggaaaacggtgtaacaagggtgaacactatcccatatcaccagctcaccgtctttcattgccatacgaaattccggatgagcattcatcaggcgggcaagaatgtgaataaaggccggataaaacttgtgcttatttttctttacggtctttaaaaaggccgtaatatccagctgaacggtctggttataggtacattgagcaactgactgaaatgcctcaaaatgttctttacgatgccattgggatatatcaacggtggtatatccagtgatttttttctccattttagcttccttagctcctgaaaatctcgataactcaaaaaatacgcccggtagtgatcttatttcattatggtgaaagttggaacctcttacgtgccgatcaacgtctcattttcgccagatatc

**pSECRETS-B:**

gacgtcttaagacccactttcacatttaagttgtttttctaatccgcatatgatcaattcaaggccgaataagaaggctggctctgcaccttggtgatcaaataattcgatagcttgtcgtaataatggcggcatactatcagtagtaggtgtttccctttcttctttagcgacttgatgctcttgatcttccaatacgcaacctaaagtaaaatgccccacagcgctgagtgcatataatgcattctctagtgaaaaaccttgttggcataaaaaggctaattgattttcgagagtttcatactgtttttctgtaggccgtgtacctaaatgtacttttgctccatcgcgatgacttagtaaagcacatctaaaacttttagcgttattacgtaaaaaatcttgccagctttccccttctaaagggcaaaagtgagtatggtgcctatctaacatctcaatggctaaggcgtcgagcaaagcccgcttattttttacatgccaatacaatgtaggctgctctacacctagcttctgGGCGAGTTTACGGGTTGTTAaaccttcgattccgacctcattaagcagctctaatgcgctgttaatcactttacttttatctaaacgagacatactcttcctttttcaatattattgaagcatttatcagggttattgtctcatgagcggatacatatttgaatgtatttagaaaaataaacaaataggggttccgcgcacatttccccgaaaagtgccacctgacgtcctctggctaactagagaacccactgcttactggcttatcgaaatttccctatcagtgatagagattgacatccctatcagtgatagagatactgagcacgtGAGACCcatgccatagcgttgttTAGGGATAACAGGGTAATaCTGTCCACACAATCTGCCCTGGTCTCcgttttagagctagaaatagcaagttaaaataaggctagtccgttatcaacttgaaaaagtggcaccgagtcggtgcttttttGGCCGGCATGGTCCCAGCCTCCTCGCTGGCGCCGGCTGGGCAACATGCTTCGGCATGGCGAATGGGACGACATCACCTCCCACAACGAagactacaccatcgttgaacagtacgaacgtgctgaaggtcgtcactccaccggtgcttaaggatccaaactcgagtaaggatctccaggcatcaaataaaacgaaaggctcagtcgaaagactgggcctttcgttttatctgttgtttgtcggtgaacgctctctactagagtcacactggctcaccttcgggtgggcctttctgcgtttatacctagggtacgggttttgctgcccgcaaacgggctgttctggtgttgctagtttgttatcagaatcgcagatccggcttcagccggtttgccggctgaaagcgctatttcttccagaattgccatgattttttccccacgggaggcgtcactggctcccgtgttgtcggcagctttgattcgataagcagcatcgcctgtttcaggctgtctatgtgtgactgttgagctgtaacaagttgtctcaggtgttcaatttcatgttctagttgctttgttttactggtttcacctgttctattaggtgttacatgctgttcatctgttacattgtcgatctgttcatggtgaacagctttgaatgcaccaaaaactcgtaaaagctctgatgtatctatcttttttacaccgttttcatctgtgcatatggacagttttccctttgatatgtaacggtgaacagttgttctacttttgtttgttagtcttgatgcttcactgatagatacaagagccataagaacctcagatccttccgtatttagccagtatgttctctagtgtggttcgttgtttttgcgtgagccatgagaacgaaccattgagatcatacttactttgcatgtcactcaaaaattttgcctcaaaactggtgagctgaatttttgcagttaaagcatcgtgtagtgtttttcttagtccgttatgtaggtaggaatctgatgtaatggttgttggtattttgtcaccattcatttttatctggttgttctcaagttcggttacgagatccatttgtctatctagttcaacttggaaaatcaacgtatcagtcgggcggcctcgcttatcaaccaccaatttcatattgctgtaagtgtttaaatctttacttattggtttcaaaacccattggttaagccttttaaactcatggtagttattttcaagcattaacatgaacttaaattcatcaaggctaatctctatatttgccttgtgagttttcttttgtgttagttcttttaataaccactcataaatcctcatagagtatttgttttcaaaagacttaacatgttccagattatattttatgaatttttttaactggaaaagataaggcaatatctcttcactaaaaactaattctaatttttcgcttgagaacttggcatagtttgtccactggaaaatctcaaagcctttaaccaaaggattcctgatttccacagttctcgtcatcagctctctggttgctttagctaatacaccataagcattttccctactgatgttcatcatctgagcgtattggttataagtgaacgataccgtccgttctttccttgtagggttttcaatcgtggggttgagtagtgccacacagcataaaattagcttggtttcatgctccgttaagtcatagcgactaatcgctagttcatttgctttgaaaacaactaattcagacatacatctcaattggtctaggtgattttaatcactataccaattgagatgggctagtcaatgataattactagtccttttcccgggtgatctgggtatctgtaaattctgctagacctttgctggaaaacttgtaaattctgctagaccctctgtaaattccgctagacctttgtgtgttttttttgtttatattcaagtggttataatttatagaataaagaaagaataaaaaaagataaaaagaatagatcccagccctgtgtataactcactactttagtcagttccgcagtattacaaaaggatgtcgcaaacgctgtttgctcctctacaaaacagaccttaaaaccctaaaggcttaagtagcaccctcgcaagctcgggcaaatcgctgaatattccttttgtctccgaccatcaggcacctgagtcgctgtctttttcgtgacattcagttcgctgcgctcacggctctggcagtgaatgggggtaaatggcactacaggcgccttttatggattcatgcaaggaaactacccataatacaagaaaagcccgtcacgggcttctcagggcgttttatggcgggtctgctatgtggtgctatctgactttttgctgttcagcagttcctgccctctgattttccagtctgaccacttcggattatcccgtgacaggtcattcagactggctaatgcacccagtaaggcagcggtatcatcaacaggcttacccgtcttactgtccctagtgcttggattctcaccaataaaaaacgcccggcggcaaccgagcgttctgaacaaatccagatggagttctgaggtcattactggatctatcaacaggagtccaagcgagctctcgaaccccagagtcccgctcagaagaactcgtcaagaaggcgatagaaggcgatgcgctgcgaatcgggagcggcgataccgtaaagcacgaggaagcggtcagcccattcgccgccaagctcttcagcaatatcacgggtagccaacgctatgtcctgatagcggtccgccacacccagccggccacagtcgatgaatccagaaaagcggccattttccaccatgatattcggcaagcaggcatcgccatgggtcacgacgagatcctcgccgtcgggcatgcgcgccttgagcctggcgaacagttcggctggcgcgagcccctgatgctcttcgtccagatcatcctgatcgacaagaccggcttccatccgagtacgtgctcgctcgatgcgatgtttcgcttggtggtcgaatgggcaggtagccggatcaagcgtatgcagccgccgcattgcatcagccatgatggatactttctcggcaggagcaaggtgagatgacaggagatcctgccccggcacttcgcccaatagcagccagtcccttcccgcttcagtgacaacgtcgagcacagctgcgcaaggaacgcccgtcgtggccagccacgatagccgcgctgcctcgtcctgcagttcattcagggcaccggacaggtcggtcttgacaaaaagaaccgggcgcccctgcgctgacagccggaacacggcggcatcagagcagccgattgtctgttgtgcccagtcatagccgaatagcctctccacccaagcggccggagaacctgcgtgcaatccatcttgttcaatcatgcgaaacgatcctcatcctgtctcttgatcagatcatgatcccctgcgccatcagatccttggcggcaagaaagccatccagtttactttgCAGGGCTTCCCAACCTTACCagagggcgccccagctggcaattcc

**pSECRETS-C (p11-LacY-wtx1)**

Available at: <http://n2t.net/addgene:69056>

tcgatgcataatgtgcctgtcaaatggacgaagcagggattctgcaaaccctatgctactccgtcaagccgtcaattgtctgattcgttaccaattatgacaacttgacggctacatcattcactttttcttcacaaccggcacggaactcgctcgggctggccccggtgcattttttaaatacccgcgagaaatagagttgatcgtcaaaaccaacattgcgaccgacggtggcgataggcatccgggtggtgctcaaaagcagcttcgcctggctgatacgttggtcctcgcgccagcttaagacgctaatccctaactgctggcggaaaagatgtgacagacgcgacggcgacaagcaaacatgctgtgcgacgctggcgatatcaaaattgctgtctgccaggtgatcgctgatgtactgacaagcctcgcgtacccgattatccatcggtggatggagcgactcgttaatcgcttccatgcgccgcagtaacaattgctcaagcagatttatcgccagcagctccgaatagcgcccttccccttgcccggcgttaatgatttgcccaaacaggtcgctgaaatgcggctggtgcgcttcatccgggcgaaagaaccccgtattggcaaatattgacggccagttaagccattcatgccagtaggcgcgcggacgaaagtaaacccactggtgataccattcgcgagcctccggatgacgaccgtagtgatgaatctctcctggcgggaacagcaaaatatcacccggtcggcaaacaaattctcgtccctgatttttcaccaccccctgaccgcgaatggtgagattgagaatataacctttcattcccagcggtcggtcgataaaaaaatcgagataaccgttggcctcaatcggcgttaaacccgccaccagatgggcattaaacgagtatcccggcagcaggggatcattttgcgcttcagccatacttttcatactcccgccattcagagaagaaaccaattgtccatattgcatcagacattgccgtcactgcgtcttttactggctcttctcgctaaccaaaccggtaaccccgcttattaaaagcattctgtaacaaagcgggaccaaagccatgacaaaaacgcgtaacaaaagtgtctataatcacggcagaaaagtccacattgattatttgcacggcgtcacactttgctatgccatagcatttttatccataagattagcggatcctacctgacgctttttatcgcaactctctactgtttctccatacccgtttttttgggctagcgattgaaaacgatgcagtttaaggtttacacctataaaagagagagccgttatcgtctgtttgtggatgtacagagtgatattattgacacgcccgggcgacggatggtgatccccctggccagtgcacgtctgctgtcagataaagtctcccgtgaactttacccggtggtgcatatcggggatgaaagctggcgcatgatgaccaccgatatggccagtgtgccggtctccgttatcggggaagaagtggctgatctcagccaccgcgaaaatgacatcaaaaacgccattaacctgatgttttggggaatataatctagcattacgctagggataacagggtaatatcacgctctagacatacggcatgcaagcttggctgttttggcggatgagagaagattttcagcctgatacagattaaatcagaacgcagaagcggtctgataaaacagaatttgcctggcggcagtagcgcggtggtcccacctgaccccatgccgaactcagaagtgaaacgccgtagcgccgatggtagtgtggggtctccccatgcgagagtagggaactgccaggcatcaaataaaacgaaaggctcagtcgaaagactgggcctttcgttttatctgttgtttgtcggtgaacgctctcctgagtaggacaaatccgccgggagcggatttgaacgttgcgaagcaacggcccggagggtggcgggcaggacgcccgccataaactgccaggcatcaaattaagcagaaggccatcctgacggatggcctttttgcgtttctacaaactcttttgtttatttttctaaatacattcaaatatgtatccgctcatgagacaataaccctgataaatgcttcaataatattgaaaaaggaagagtatgagtattcaacatttccgtgtcgcccttattcccttttttgcggcattttgccttcctgtttttgctcacccagaaacgctggtgaaagtaaaagatgctgaagatcagttgggtgcacgagtgggttacatcgaactggatctcaacagcggtaagatccttgagagttttcgccccgaagaacgttttccaatgatgagcacttttaaagttctgctatgtggcgcggtattatcccgtgttgacgccgggcaagagcaactcggtcgccgcatacactattctcagaatgacttggttgagtactcaccagtcacagaaaagcatcttacggatggcatgacagtaagagaattatgcagtgctgccataaccatgagtgataacactgcggccaacttacttctgacaacgatcggaggaccgaaggagctaaccgcttttttgcacaacatgggggatcatgtaactcgccttgatcgttgggaaccggagctgaatgaagccataccaaacgacgagcgtgacaccacgatgcctgcagcaatggcaacaacgttgcgcaaactattaactggcgaactacttactctagcttcccggcaacaattaatagactggatggaggcggataaagttgcaggaccacttctgcgctcggcccttccggctggctggtttattgctgataaatctggagccggtgagcgtgggtctcgcggtatcattgcagcactggggccagatggtaagccctcccgtatcgtagttatctacacgacggggagtcaggcaactatggatgaacgaaatagacagatcgctgagataggtgcctcactgattaagcattggtaactgtcagaccaagtttactcatatatactttagattgatttacgcgccctgtagcggcgcattaagcgcggcgggtgtggtggttacgcgcagcgtgaccgctacacttgccagcgccctagcgcccgctcctttcgctttcttcccttcctttctcgccacgttcgccggctttccccgtcaagctctaaatcgggggctccctttagggttccgatttagtgctttacggcacctcgaccccaaaaaacttgatttgggtgatggttcacgtagtgggccatcgccctgatagacggtttttcgccctttgacgttggagtccacgttctttaatagtggactcttgttccaaacttgaacaacactcaaccctatctcgggctattcttttgatttataagggattttgccgatttcggcctattggttaaaaaatgagctgatttaacaaaaatttaacgcgaattttaacaaaatattaacgtttacaatttaaaaggatctaggtgaagatcctttttgataatctcatgaccaaaatcccttaacgtgagttttcgttccactgagcgtcagaccccgtagaaaagatcaaaggatcttcttgagatcctttttttctgcgcgtaatctgctgcttgcaaacaaaaaaaccaccgctaccagcggtggtttgtttgccggatcaagagctaccaactctttttccgaaggtaactggcttcagcagagcgcagataccaaatactgtccttctagtgtagccgtagttaggccaccacttcaagaactctgtagcaccgcctacatacctcgctctgctaatcctgttaccagtggctgctgccagtggcgataagtcgtgtcttaccgggttggactcaagacgatagttaccggataaggcgcagcggtcgggctgaacggggggttcgtgcacacagcccagcttggagcgaacgacctacaccgaactgagatacctacagcgtgagctatgagaaagcgccacgcttcccgaagggagaaaggcggacaggtatccggtaagcggcagggtcggaacaggagagcgcacgagggagcttccagggggaaacgcctggtatctttatagtcctgtcgggtttcgccacctctgacttgagcgtcgatttttgtgatgctcgtcaggggggcggagcctatggaaaaacgccagcaacgcggcctttttacggttcctggccttttgctggccttttgctcacatgttctttcctgcgttatcccctgattctgtggataaccgtattaccgcctttgagtgagctgataccgctcgccgcagccgaacgaccgagcgcagcgagtcagtgagcgaggaagcggaagagcgcctgatgcggtattttctccttacgcatctgtgcggtatttcacaccgcatacgtacgatttaaataggcctgactcactatagggagaccggaattccctggcacgacaggtttcccgactggaaagcgggcagtgagcgcaacgcaattaatgtgagttagctcactcattagggaccccgggctttacactttatgcttccggctcgtatgttgtgtggaattgtgagcggataacaatttcacacaggaaacagctatgaccatgattacggattcactggccgtcgttttacaacgtcgtgactggtaaaacccgggcgttacccaacttaatcgccttgcagcacatccccctttcgccagcaggcgtaataaggaaaggatccatgtactatttgaaaaacacaaacttttggatgttcggtttattctttttcttttacttttttatcatgggagcctacttcccgtttttcccgatttggctacatgatatcaaccatatcagcaaaagtgatacgggtattatttttgccgctatttctctgttctcgctattattccaaccgctgtttggtctgctttctgacaaactcggtctacgcaaatacctgctgtggattattaccggcatgttagtgatgtttgcgccgttctttatttttatcttcgggccactgctgcagtacaacattttagtaggatcgattgttggtggtatttatctaggctttagttttaacgccggtgcgccagcagtagaggcatttattgagaaagtcagccggcgcagtaatttcgaatttggtcgcgcgcggatgtttggcagtgttggctgggcgctggttgcctcgattgtcgggatcatgttcaccattaataatcagtttgttttctggctgggctctggcagttgtctcatcctcgccgttttactctttttcgccaaaacggacgcgccctcgagtgccacggttgccaatgcggtaggtgccaaccattcggcatttagccttaagctggcactggaactgttcagacagccaaaactgtggtttttgtcactgtatgttattggcgtttcctccacctacgatgtttttgaccaacagtttgctaatttctttacttcgttctttgctaccggtgaacagggtacccgcgtatttggctacgtaacgacaatgggcgaattacttaacgcctcgattatgttctttgcgccactgatcattaatcgcatcggtgggaagaatgccctgctgctggctggcactattatgtctgtacgtattattggctcatcgttcgccacctcagcgctggaagtggttattctgaaaacgctgcatatgtttgaagtaccgttcctgctggtgggctcctttaaatatattactagtcagtttgaagtgcgtttttcagcgacgatttatctggtcagtttcagcttctttaagcaactggcgatgatttttatgtctgtactggcgggcaatatgtatgaaagcataggtttccaaggcgcttatctggtgctgggtctggtggcgctgggcttcaccttaatttccgtgttcacgcttagcggcccgggcccgctttccctgctgcgtcgtcaggtgaatgaagtcgcttaaaggcc
